# COMBO-RATE: An experimentally validated bioinformatic tool to identify promiscuous HLA restrictions

**DOI:** 10.64898/2026.04.14.718521

**Authors:** Jessica Nevarez-Mejia, Raphael Trevizani, Adam Abawi, Emil Johansson, Aaron Sutherland, Alba Grifoni, Ricardo da Silva Antunes, Alessandro Sette

**Affiliations:** Center for Vaccine Innovation, La Jolla Institute for Immunology (LJI), La Jolla, CA 92037, USA; Fiocruz Ceará, R. São José, S/N - Precabura, Eusébio - CE, 61773-270, Brazil; Department of Medicine, Division of Infectious Diseases and Global Public Health, University of California, San Diego (UCSD), La Jolla, CA 92037, USA

## Abstract

Defining HLA restriction of T cell epitopes is essential for understanding immune responses in infectious disease, autoimmunity, and vaccine design. Current bioinformatic programs, including the IEDB RATE tool, enable inference of single-HLA restrictions from immune response data of HLA-typed donors. However, T cell epitopes are frequently presented by multiple HLA alleles, a phenomenon termed promiscuous restriction, limiting the utility of single-allele approaches. To address this limitation, we developed COMBO-RATE, an extension of RATE that systematically evaluates combinations of HLA alleles to identify multi-allelic restriction patterns. Analysis of three independent datasets spanning distinct antigen systems and different epitope discovery strategies revealed that promiscuous restriction is a near-universal feature of immunodominant epitopes. Focusing on the 43 immunodominant CD4⁺ T cell epitopes identified in a *B. pertussis* genome-wide screen, COMBO-RATE outperformed conventional RATE, identifying restrictions for 35 of 43 epitopes, compared to 24 by RATE alone, and uncovered 64 additional allele restrictions, including 29 unique alleles. Experimental validation using single-HLA transfected cell lines and antigen presentation assays confirmed COMBO-RATE-inferred restrictions, demonstrating that a single epitope can be independently presented by distinct HLA alleles. Overall, COMBO-RATE provides a robust and scalable framework for defining complete HLA restriction profiles from existing population response data, with important implications for the design of vaccines requiring broad HLA coverage across genetically diverse populations. This pipeline is available as both a Python package and a user-friendly web application.

## INTRODUCTION

T cells express specific T Cell Receptors (TCRs) which recognize a complex between a peptide (termed epitope), and MHC molecules (HLA in humans)^1^. The HLA restriction of a given epitope indicates the specific HLA allelic variant(s) that bind and present that epitope for T cell recognition^2^. Knowledge of HLA restriction of T cell epitopes is important for several immunological applications, to enable production of multimeric/tetrameric staining reagents^3^, to select donors expected to respond to a given peptide in longitudinal investigation^4^, and to project population coverage afforded by different epitope combinations^5^. This is of particular importance for the design of vaccines that provide broad population coverage and ensure effectiveness across different ethnic populations and geographic regions with diverse HLA types^6^.

Several different studies have further shown different peptide binding of different HLA molecules can be largely overlapping, which as results can be grouped in broad HLA supertypes^7,8^. As a result, a given epitope can bind to multiple HLA allelic variants, a phenomenon termed “degenerate binding”^9^. Epitopes associated with degenerate binding can be recognized in the context of multiple HLA variants and thus be associated with multiple HLA restrictions^10^. These epitopes are commonly referred to as “promiscuous” epitopes^11^. This is particularly relevant, since promiscuous epitopes tend to be dominant and account for a large fraction of the response^12^.

HLA restriction of a given epitope can be experimentally determined in antigen presentation assays using panels of cell lines expressing defined HLA molecules^13^, but these assays are laborious and alternative computational methods have been utilized. These methodologies rely on predicting the peptide’s binding capacity for the HLA molecules expressed in the responding individuals^14^ and determining whether any particular HLA molecule(s) is associated with peptide responsiveness^15^. In particular, the RATE program, available in the IEDB^15,16^, determines likely potential restriction based on known response patterns of group of HLA typed individuals. This approach is particularly valuable for prioritizing which HLA alleles to test experimentally, thereby saving time and resources in the characterization process of immune responses to a pathogen or disease.

However, no well-defined and automated pipeline is available for determining promiscuous restrictions, potentially underestimating the true breadth of an epitope’s restriction. Here we modified and expanded the RATE approach to pinpoint potential promiscuous restriction and experimentally validated this approach in selected examples.

## RESULTS

### Promiscuous restrictions are common amongst dominant epitopes

In the context of a genome-wide screening of *Bordetella Pertussis* (BP) human T cell epitopes^17^, we identified a panel of 43 HLA class II epitopes, that were dominantly recognized (defined as recognized by > 4 donors). These epitopes were further tested in an additional cohort of 20 donors, resulting in testing an average of 34 HLA-typed donors per epitope (range 20-46 donors) (Nevarez-Mejia et al., manuscript in preparation). The pattern of responsiveness to each peptide for each donor (as a binary yes/no outcome) and the HLA types of the individuals tested are provided as **Table S1** and **S2**, respectively. As a first step we quantitated how frequently promiscuous restrictions were observed. We reasoned that by definition, the HLA restricting molecule(s) must be expressed in the responding individual. If a single HLA restriction is associated with the epitope, that HLA molecules must be expressed in all responders. If no single HLA molecules is found in every single responding individual, then, also by definition, that epitope is restricted by multiple HLAs (promiscuous recognition).

To this end, for each epitope, we tabulated the HLA class II types expressed in individuals associated with positive responses and recorded the most frequent HLA class II molecule(s) expressed in the responding donors. The results are shown in **Figure 1** where for each epitope the frequency of the HLA class II molecule most frequently expressed in the responding individuals is plotted. In only three cases out of 43 epitopes, a single HLA variant was common amongst all responders. Specifically, HLA DRB4*01:01 was found in 6/6 responders for peptide 12, HLA DPA1*01:03/B1*04:01 in 6/6 responders for peptide 33 and HLA DPA1*01:03/B1*02:01 was found in 3/3 responders for peptide 41. In all other cases, no single HLA could explain all responses (**Figure 1A**). The median frequency of the HLA class II molecule most frequently expressed in the responding individuals was 0.60 (range 0.46 to 1), and the frequency of the most common HLA correlated inversely with the number of responder donors (R=-0.41, *p*=0.007), further suggesting that frequently recognized donors are not restricted to a single HLA (**Figure 1B**). This data underlines how multiple HLA restrictions are commonly associated with dominant epitopes, at least in the BP dataset.

**Figure 1.**
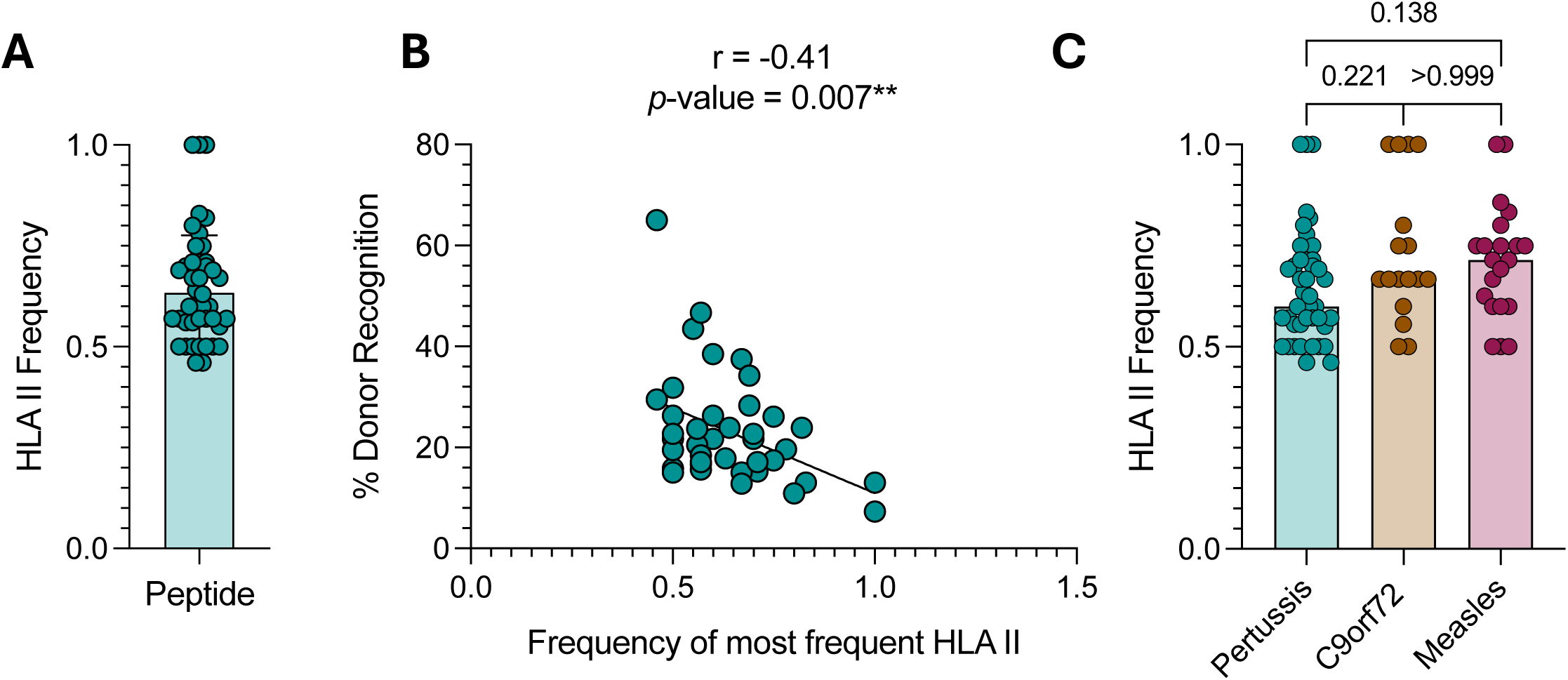
Promiscuous HLA restriction is frequently observed among dominant epitopes. **A)** Frequency of the HLA class II molecule most frequently expressed in the responding individuals for each peptide. **B**) Correlation between the frequency of HLAs with the % of donor recognition. Spearman correlation was performed. **C)** Proportion of the most frequent HLA II among positive donors between pertussis, measles, and ALS datasets. Frequency of the HLA class II molecule most frequently expressed in the responding individuals for each peptide between predicted (Pertussis) and non- predicted epitope (C9orf72 and Measles) screening studies. Statistical significance was determined using Krustal-Wallis test followed by Dunn’s multiple comparison test.

### Promiscuous HLA restriction patterns are independent of peptide selection strategy

The fact that multiple HLA restrictions are associated with the BP data set was not entirely surprising, because the BP peptide dataset comprised dominant CD4+ T cell epitopes identified using promiscuous HLA class II binding predictions^17,18^. To confirm that the observed promiscuous restrictions were not limited to this prediction-based selection approach, we performed an analysis similar to **Figure 1A** on CD4+ datasets from recent epitope identification studies that used unbiased peptide selection. Specifically, we examined HLA class II types associated with positive responses in studies that screened unbiased overlapping peptides spanning entire open reading frames (ORFs) of the Measles virus proteome (under review) and the human C9orf72 protein^19^, rather than computationally predicted epitopes.

**Figure 1C** shows the frequency of the most commonly expressed HLA class II molecule among responders for each epitope for these two datasets. While the median values are marginally higher in the Measles (median 0.71, range 0.50 – 1.00) and C9orf72 (median 0.67, range 0.50 – 1.00) datasets, no significant difference is observed, and a single HLA molecule could potentially be a restriction element for only 2-4 epitopes in the different datasets. In summary, this data indicates that most epitopes are associated with promiscuous restrictions, in three different data sets, and independently of the epitope identification strategy utilized.

### A strategy to assess potential multiple HLA restrictions

The RATE program^15,16^ was developed to infer restrictions based on epitope response patterns in HLA typed individuals, by identifying HLA alleles more frequently expressed in responders as compared to non-responders. Specifically, RATE accepts as input the patterns of responses to one or more epitopes in a set of given individuals, and a separate file detailing the HLA types expressed in those individuals. The program returns as output, for each epitope and for each HLA expressed in the tested population, the number of individuals in four categories: HLA-positive responders (A+R+), HLA-positive non-responders (A+R-), HLA-negative responders (A-R+), and HLA-negative non-responders (A-R-). Relative frequency (RF) and odds ratios (OR) are calculated, with statistical significance determined by Fisher’s exact test (without Bonferroni correction).

Here, we built on this algorithm and developed an original workflow, as exemplified by a sample output for epitope 23 (P23) shown in **Table 1 and Table 2**. The HLA types of ten individuals responding to this epitope are shown in **Table S3.** Overall, 35 different HLA molecules are expressed in the 10 responder individuals (**Table 1**; “Alleles”). We further considered only the 9 HLAs predicted to bind the epitope with a 25%-ile value or better (**Table 1**; “25%-ile rank”). We next identified the p-values provided by the RATE program for the HLA alleles, only considering positive associations (RF≥1.0) (**Table 1**; “RATE p-value”). In this case it was found that DRB1*15:01 was associated with a statistically significant p-value of 0.048. However, DRB1*15:01 was present only in 5 out of 10 responding individuals, which indicates the presence of additional restricting HLAs for P23 (**Table 2**).

**Table 1:** Sample workflow for epitope P23. P23 binding capacity of the 35 different HLA molecules that are expressed in the 10 responder individuals and RATE p-values for alleles predicted to bind P23 at the top 25%-ile rank.

| Alleles | 25%-ile rank | RATE p-value |
| --- | --- | --- |
| DRB1*11:01 | 4.4 | 0.539 |
| DRB1*01:01 | 12 | 0.113 |
| DRB1*08:02 | 14 | 0.222 |
| DRB1*13:01 | 15 | 1 |
| DRB1*15:01 | 18 | 0.048 |
| DQA1*01:01/DQB1*05:01 | 20 | 0.349 |
| DRB1*07:01 | 22 | 1 |
| DRB4*01:01 | 23 | 0.481 |
| DRB1*14:02 | 24 | 0.222 |
| DPA1*01:03/DPB1*06:01 | 27 | - |
| DRB5*01:02 | 28 | - |
| DPA1*01:03/DPB1*04:01 | 31 | - |
| DRB1*15:02 | 31 | - |
| DRB3*02:02 | 32 | - |
| DRB5*01:01 | 33 | - |
| DRB1*04:03 | 34 | - |
| DRB3*01:01 | 35 | - |
| DPA1*01:03/DPB1*02:01 | 35 | - |
| DPA1*01:03/DPB1*20:01 | 38 | - |
| DPA1*01:03/DPB1*16:01 | 38 | - |
| DPA1*01:03/DPB1*03:01 | 41 | - |
| DQA1*05:01/DQB1*02:01 | 41 | - |
| DRB1*04:01 | 45 | - |
| DRB1*03:01 | 48 | - |
| DPA1*01:03/DPB1*04:02 | 49 | - |
| DQA1*04:01/DQB1*04:02 | 53 | - |
| DQA1*02:01/DQB1*02:02 | 55 | - |
| DQA1*03:01/DQB1*03:02 | 57 | - |
| DPA1*02:01/DPB1*01:01 | 59 | - |
| DQA1*05:01/DQB1*03:01 | 69 | - |
| DQA1*05:01/DQB1*06:02 | 73 | - |
| DQA1*01:03/DQB1*06:03 | 82 | - |
| DQA1*01:02/DQB1*06:02 | 82 | - |
| DQA1*03:01/DQB1*03:01 | 83 | - |
| DQA1*01:03/DQB1*06:01 | 84 | - |

**Table 2:** COMBO-RATE results in RATE format for P23.

| HLA | Peptide | A+R+ | A-R+ | A+R- | A-R- | N_donors | Relative_<br>freq | Odds_<br>ratio | Fisher_<br>pval |
| --- | --- | --- | --- | --- | --- | --- | --- | --- | --- |
| DRB1*15:01 | TRFSNKIRLMGPLQV | 5 | 5 | 6 | 29 | 45 | 2.05 | 5 | 0.048 |
| DRB1*15:01+DRB1*01:01 | TRFSNKIRLMGPLQV | 8 | 2 | 8 | 28 | 46 | 2.30 | 14 | 0.0015 |
| DRB1*15:01+DRB1*01:01+<br>DRB1*08:02+DRB1*14:02 | TRFSNKIRLMGPLQV | 9 | 1 | 8 | 28 | 46 | 2.44 | 32 | 0.00018 |
| DRB1*15:01+DRB1*01:01+<br>DRB1*14:02+DRB1*08:02+<br>DQA1*01:01/DQB1*05:01 | TRFSNKIRLMGPLQV | 9 | 1 | 10 | 26 | 46 | 2.18 | 23 | 0.00063 |

A simple issue limits the power of the single inferred restrictions, namely the fact that if multiple restrictions are present, the positive donors for one restriction will confound and decrease the significance of the association with the other restrictions, which might be detected just as positive but not significant trends. To overcome this limitation we decided to record, for any given peptide, the alleles associated with those positive trends and run associations with epitope responsiveness for combinations of these alleles. We denominated this approach as “COMBO-RATE”.

### COMBO-RATE identifies potential multiple HLA restrictions

The algorithm employed in COMBO-RATE starts by combining the two HLAs associated with the lowest p-values (significant or trends) and records the new p-value. If the p-value decreases, then the effect of further combining the HLA allele with the third-lowest p-value is evaluated, and so on. The process is repeated until adding additional HLAs no longer reduces the obtained p-value.

In the case of the P23 epitope, there is a trend for DRB1*01:01 (p-value =0.113); combining DRB1*15:01 and DRB1*01:01 yields a lower significant p-value of 0.0015. Further combination with DRB1*08:02 and DRB1*14:02 (which showed equivalent p-values) yielded the most significant association (p=0.0002) (**Table 2**). Adding further HLA variants did not further improve p-values. The combination of these alleles is now associated with 9 restriction-matched donors out of the 10 responders.

The results for the whole set of BP epitopes are shown in **Table 3** (left side) for the existing RATE program. Overall significant restrictions could be inferred for 24 of the 43 epitopes, and these inferred restrictions only account in average for 35% of the total donor/peptide instances (139/401). The results for the COMBO-RATE approach are also summarized in **Table 3** (right side) in the columns labeled as “Newly identified COMBO-RATE restrictions” along with the percentage of allele-assigned donors. For 17/43 peptides the COMBO-RATE approach assigns additional restriction-matched donors (p<0.0001) (**Figure 2A**). Importantly, COMBO-RATE further infers restrictions in 8 cases where no restriction was inferred with individual alleles. Overall, the COMBO- RATE grouping inferred restrictions for 220/401 (55%) donor/peptide instances, a significant increase (p<0.0001) from the conventional RATE method (139/401, 35%) (**Figure 2B**), and yielded markedly improved p-values for the inferred restrictions (**Figure 2C**). If we exclude the peptides for which no restriction is available, the number is 220/330 (67%). In addition, COMBO-RATE identifies in this epitope data set an overall gain of 64 new allele restrictions, with 29 representing unique alleles.

**Figure 2:**
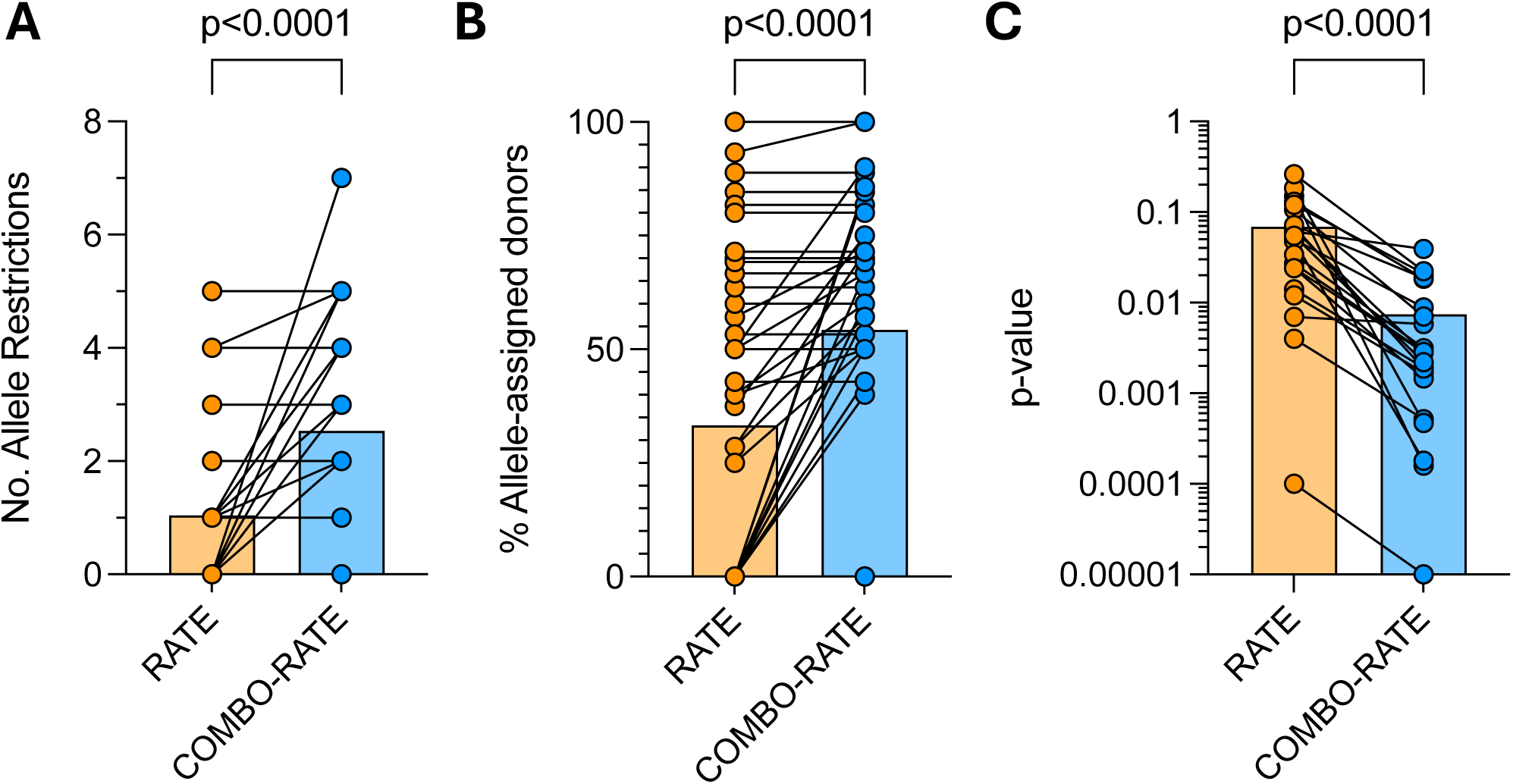
COMBO-RATE identified new epitope restrictions and significantly increases the percentage of allele-matched donors. **A)** Number of identified allele restrictions between RATE and COMBO-RATE (p<0.0001). **B)** % allele-assigned donors between RATE and COMBO-RATE (p<0.0001). **C**) p-value of inferred restrictions between RATE and COMBO-RATE (p<0.0001). Data graphed in log scale. Statistical significance was determined by Wilcoxon matched-pairs signed rank test.

**Table 3:** Identified RATE and COMBO-RATE restrictions for all epitopes.

| Peptide | RATE restrictions | No. of RATE restrictions | RATE allele-assigned donors | Newly identified COMBO-RATE Restrictions | No. Restrictions gained with COMBO-RATE | COMBO-RATE allele-assigned donors | Total No. of Restrictions | % Gained in coverage of explained donors |
| --- | --- | --- | --- | --- | --- | --- | --- | --- |
| 1 | DRB1*07:01;DRB1*04:07;DRB1*04:04;DQA1*02:01/DQB1*02:02;DQA1*03:01/DQB1*03:02 | 5 | 8/9 (89%) |  | 0 | 8/9 (89%) | 5 | 0% |
| 2 | DRB1*07:01 | 1 | 5/10 (50%) |  | 0 | 5/10 (50%) | 1 | 0% |
| 3 |  | 0 | 0/12 (0%) |  | 0 | 0/12 (0%) | 0 | 0% |
| 4 | DRB1*07:01 | 1 | 8/20 (40%) | DRB1*04:07 | 1 | 10/20 (50%) | 2 | 10% |
| 5 | DRB1*04:04;DQA1*02:01/DQB1*02:02;DPA1*02:01/DPB1*01:01;DRB4*01:01 | 4 | 11/13 (85%) |  | 0 | 11/13 (85%) | 4 | 0% |
| 6 | DRB1*04:07 | 1 | 3/8 (38%) | DPA1*01:03/DPB1*14:01;DRB1*04:04;DRB1*04:01;DQA1*03:01/DQB1*03:01 | 4 | 6/8 (75%) | 5 | 38% |
| 7 | DRB4*01:01;DRB1*07:01 | 2 | 9/11 (82%) |  | 0 | 9/11 (82%) | 2 | 0% |
| 8 |  | 0 | 0/14 (0%) |  | 0 | 0/14 (0%) | 0 | 0% |
| 9 |  | 0 | 0/13 (0%) |  | 0 | 0/13 (0%) | 0 | 0% |
| 10 | DRB1*04:01 | 1 | 4/10 (40%) | DRB1*11:01;DRB1*11:04 | 2 | 6/10 (60%) | 3 | 20% |
| 11 | DQA1*03:01/DQB1*03:02 | 1 | 4/7 (57%) | DRB1*04:07 | 1 | 5/7 (71%) | 2 | 14% |
| 12 | DRB4*01:01 | 1 | 6/6 (100%) | DRB1*04:02;DQA1*03:01/DQB1*03:01 | 2 | 6/6 (100%) | 3 | 0% |
| 13 | DRB1*14:05;DRB1*15:01;DRB5*01:01;DQA1*01:02/DQB1*06:02 | 4 | 6/10 (60%) |  | 0 | 6/10 (60%) | 4 | 0% |
| 14 | DQA1*02:01/DQB1*02:02;DRB1*04:02;DRB1*07:01 | 3 | 7/10 (70%) |  | 0 | 7/10 (70%) | 3 | 0% |
| 15 | DRB1*03:01;DRB1*12:01;DQA1*05:01/DQB1*02:01 | 3 | 8/15 (53%) |  | 0 | 8/15 (53%) | 3 | 0% |
| 16 | DPA1*01:03/DPB1*04:02;DQA1*01:01/DQB1*05:01;DRB1*11:04;DRB1*01:01 | 4 | 14/15 (93%) | DRB1*14:05 | 1 | 15/15 (100%) | 5 | 7% |
| 17 | DRB5*01:01;DRB1*15:01;DQA1*01:02/DQB1*06:02 | 3 | 7/11 (64%) |  | 0 | 7/11 (64%) | 3 | 0% |
| 18 |  | 0 | 0/10 (0%) | DRB1*11:01;DPA1*02:01/DPB1*01:01 | 2 | 4/10 (40%) | 2 | 40% |
| 19 |  | 0 | 0/6 (0%) |  | 0 | 0/6 (0%) | 0 | 0% |
| 20 | DRB1*04:04 | 1 | 2/7 (29%) | DQA1*05:01/DQB1*02:02;DRB5*01:02;DRB1*07:01;DRB1*15:02 | 4 | 5/7 (71%) | 5 | 43% |
| 21 | DPA1*01:03/DPB1*11:01 | 1 | 2/7 (29%) | DRB1*15:02 | 1 | 4/7 (57%) | 2 | 29% |
| 22 |  | 0 | 0/6 (0%) | DRB1*14:02;DPA1*02:02/DPB1*05:01;DRB1*01:03;DRB1*04:05 | 4 | 4/6 (67%) | 4 | 67% |
| 23 | DRB1*15:01 | 1 | 5/10 (50%) | DRB1*01:01;DRB1*14:02;DRB1*08:02 | 3 | 9/10 (90%) | 4 | 40% |
| 24 |  | 0 | 0/9 (0%) | DRB1*01:01;DRB5*01:02;DQA1*03:01/DQB1*03:03;DRB1*09:01;DQA1*06:01/DQB1*03:01 | 5 | 6/9 (67%) | 5 | 67% |
| 25 |  | 0 | 0/7 (0%) | DRB1*14:04;DPA1*02:02/DPB1*05:01;DRB1*13:02;DRB1*04:02;DRB1*08:03 | 5 | 4/7 (57%) | 5 | 57% |
| 26 | DPA1*01:03/DPB1*11:01 | 1 | 2/8 (25%) | DRB1*12:01 | 1 | 4/8 (50%) | 2 | 25% |
| 27 | DQA1*03:01/DQB1*03:02 | 1 | 4/6 (67%) |  | 0 | 4/6 (67%) | 1 | 0% |
| 28 |  | 0 | 0/10 (0%) | DPA1*01:03/DPB1*06:01;DPA1*01:03/DPB1*05:01;DRB1*15:02 | 3 | 6/10 (60%) | 3 | 60% |
| 29 |  | 0 | 0/10 (0%) | DRB1*04:01;DRB1*04:07;DRB1*04:05 | 3 | 5/10 (50%) | 3 | 50% |
| 30 |  | 0 | 0/6 (0%) |  | 0 | 0/6 (0%) | 0 | 0% |
| 31 |  | 0 | 0/9 (0%) |  | 0 | 0/9 (0%) | 0 | 0% |
| 32 | DRB5*01:01;DRB1*15:01 | 2 | 4/5 (80%) |  | 0 | 4/5 (80%) | 2 | 0% |
| 33 | DRB1*03:01 | 1 | 3/6 (50%) | DRB1*14:02;DQA1*02:01/DQB1*02:01;DQA1*05:01/DQB1*06:02 | 3 | 4/6 (67%) | 4 | 17% |
| 34 | DRB3*02:02 | 1 | 9/13 (69%) |  | 0 | 9/13 (69%) | 1 | 0% |
| 35 |  | 0 | 0/6 (0%) | DRB1*11:01;DPA1*01:03/DPB1*09:01 | 2 | 3/6 (50%) | 2 | 50% |
| 36 |  | 0 | 0/7 (0%) | DRB1*04:04;DRB1*14:05;DRB1*04:03;DRB4*01:03 | 4 | 3/7 (43%) | 4 | 43% |
| 37 |  | 0 | 0/8 (0%) |  | 0 | 0/8 (0%) | 0 | 0% |
| 38 |  | 0 | 0/14 (0%) | DRB1*03:01;DRB3*01:01;DRB1*12:01 | 3 | 8/14 (57%) | 3 | 57% |
| 39 |  | 0 | 0/13 (0%) | DQA1*04:01/DQB1*04:02;DQA1*03:01/DQB1*03:02;DQA1*01:02/DQB1*06:02;DQA1*01:03/DQB1*06:03;DQA1*03:01/DQB1*03:01;DPA1*01:03/DPB1*17:01;DQA1*05:01/DQB1*03:01 | 7 | 11/13 (85%) | 7 | 85% |
| 40 | DPA1*01:03/DPB1*02:01 | 1 | 5/7 (71%) |  | 0 | 5/7 (71%) | 1 | 0% |
| 41 |  | 0 | 0/3 (0%) |  | 0 | 0/3 (0%) | 0 | 0% |
| 42 |  | 0 | 0/7 (0%) | DRB1*03:01;DRB1*01:03;DRB1*07:01 | 3 | 6/7 (86%) | 3 | 86% |
| 43 | DQA1*05:01/DQB1*02:01 | 1 | 3/7 (43%) |  | 0 | 3/7 (43%) | 1 | 0% |

In summary, HLA restrictions could now be inferred for 35 of the 43 of the epitopes. These results underline how the new COMBO-RATE strategy greatly improves the number and significance of inferred restrictions.

### Experimental validation of new COMBO-RATE inferred HLA restrictions

To experimentally validate the restrictions inferred by the COMBO-RATE approach, we selected P23 as a representative example, along with donors with known response profiles, as this epitope had previously been used to illustrate the COMBO- RATE algorithm (**Table 1 and Table 2**). As a proof of concept, The RATE-inferred restriction (DRB1*15:01) and the COMBO-RATE-inferred restriction (DRB1*01:01) for P23 were experimentally validated using cell lines individually transfected with single HLA class II molecules, in an antigen presentation assay^13^. To isolate the contribution of each allele independently, RATE-inferred restrictions were tested in donors expressing DRB1*15:01 but not DRB1*01:01 (n=3), and COMBO-RATE-inferred restrictions were tested in donors expressing DRB1*01:01 but not DRB1*15:01 (n=2). In both groups, donors lacked DRB1*08:02 and DRB1*14:02 expression, additional COMBO-RATE- inferred restrictions, to minimize confounding contributions from other HLA-DR alleles (**Table S3**).

Single HLA class II transfected cell lines expressing HLA-DR molecules (L466.1 (DRB1*15:01) and L554.3 (DRB1*01:01)) were used as antigen-presenting cells (APCs) (**Sup. Figure 1**). The parental cell line lacking HLA-DR expression (DAP3) was used as a negative control. Briefly, donor PBMCs were stimulated with P23 to expand antigen-specific T cells. After two weeks of *in vitro* expansion, epitope-specific T cells were co-cultured with P23-pulsed or unpulsed APCs. T cell activation was assessed using FluoroSpot assay measuring the release of the Th1 pro-inflammatory cytokine interferon-γ (IFN-γ)^13^. Donors expressing the RATE-associated DRB1*15:01 allele (n=3) exhibited higher levels of IFN-γ producing T cells following co-culture with P23-pulsed L466.1 (DRB1*15:01) APCs compared to P23-pulsed L554.3 (DRB1*01:01) APCs and compared to the parental HLA-DR-negative cell line (mean net response = 3273 spot forming cells (SFC)) (**Figure 3A**). Consistently, these donors showed higher IFN-γ responses to pulsed L466.1 APCs compared to unpulsed L466.1 controls (**Figure 3B, Sup. Figure 2A**). As a positive control for epitope specificity, donor PBMCs stimulated with P23 in the absence of APCs also showed high levels of IFN-γ compared to DMSO-treated controls (**Sup. Figure 2A**).

**Figure 3.**
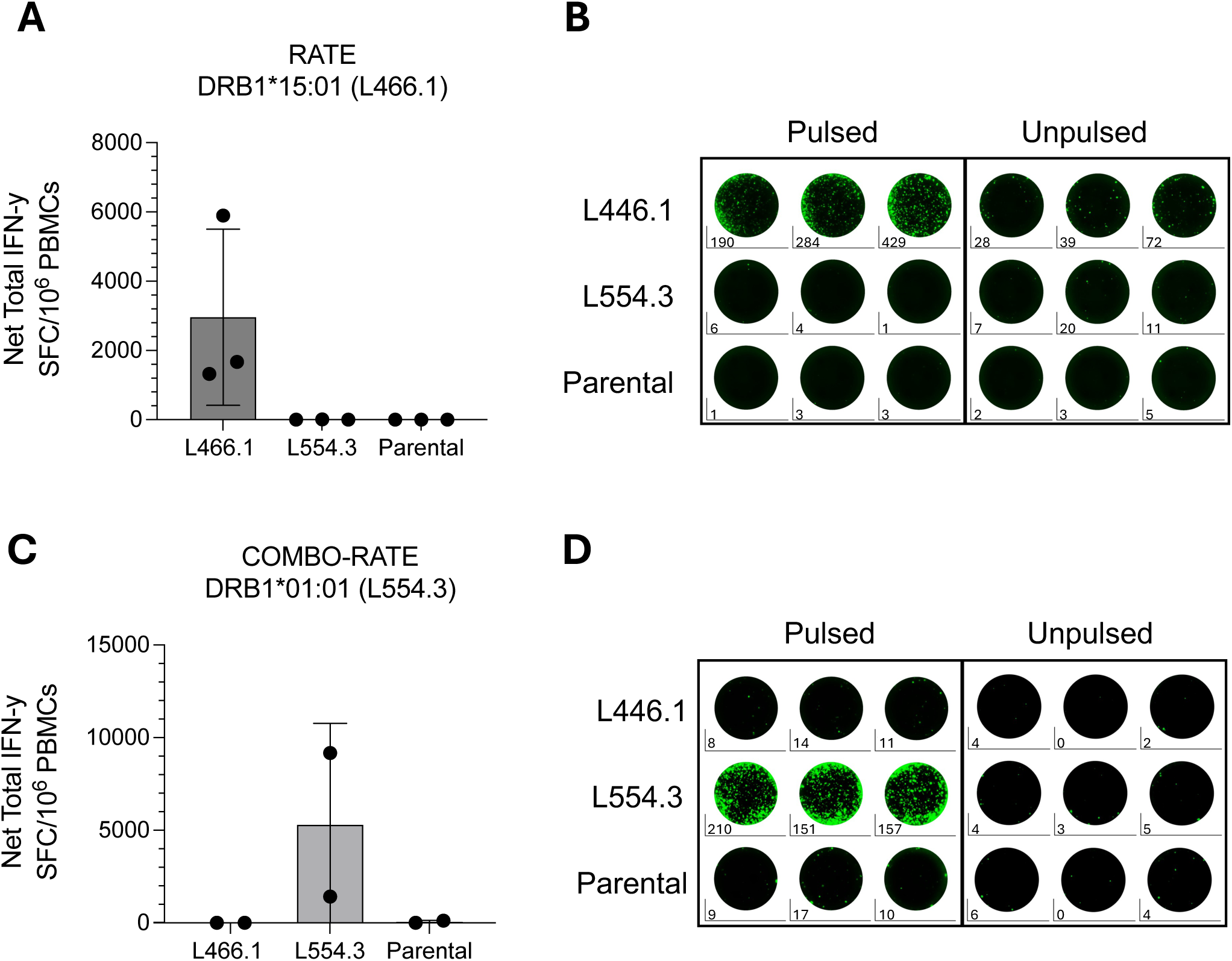
Experimental validation of RATE and COMBO-RATE restrictions. **A)** Net total IFN-y levels (spot forming cell (SFC)/10^6^ PBMCs) of RATE-associated DRB1*15:01 allele donors (n=3) when co-cultured with P23-pulsed L466.1 (DRB1*15:01), L554.3 (DRB1*01:01), and parental antigen presenting cells (APCs). Net total denotes signal of P23 pulsed minus unpulsed. **B)** Representative FluoroSpot images illustrating triplicate wells for one RATE-associated DRB1*15:01 allele donor when co-cultured with P23- pulsed and unpulsed APC lines. **C)** Net total IFN-y levels (SFC/10^6^ PBMCs) of COMBO- RATE-associated DRB1*15:01 allele donors (n=2) when co-cultured with P23-pulsed L466.1 (DRB1*15:01), L554.3 (DRB1*01:01), and parental APCs. Net total denotes signal of P23 pulsed minus unpulsed. **D)** Representative FluoroSpot images illustrating triplicate wells for one COMBO-RATE-associated DRB1*01:01 donor when co-cultured with P23-pulsed and unpulsed APC lines.

Similarly, donors expressing the COMBO-RATE-inferred DRB1*01:01 allele (n=2) exhibited higher levels of IFN-γ producing T cells following co-culture with P23-pulsed L554.3 (DRB1*01:01) APCs compared to P23-pulsed L466.1 (DRB1*15:01) APCs and compared to the parental HLA-DR-negative cell line (mean net response = 6777 SFC) (**Figure 3C**). Consistently, these donors showed higher IFN-γ responses to pulsed L554.3 APCs compared to unpulsed L554.3 controls (**Figure 3D, Sup. Figure 2B**). Stimulation of donor PBMCs with P23 in the absence of APCs further confirmed that the observed responses were epitope-specific (**Sup. Figure 2B**). In summary, these findings demonstrate that the two distinct HLA-DR alleles DRB1*15:01 and DRB1*01:01 can independently present the same peptide to CD4⁺ T cells, confirming the promiscuous restriction of P23 and validating COMBO-RATE as a tool for identifying additional HLA restrictions beyond those detected by single-allele analysis.

### A web-phase interface to make COMBO-RATE available to the scientific community

To make COMBO-RATE widely available to the scientific community, we created a web interface (https://comborate.onrender.com), which implements the standalone Python script available at: https://github.com/raphaeltrevizani/comborate_shared/tree/master.

COMBO-RATE extends the functionality of the RATE^15,16^ program that iteratively combines different HLA alleles into supertypes and reruns the RATE analysis based on the supertypes so long as significance of the Fisher p-values improve. Mandatory input data includes (I) the HLA-typed donors (**Figure 4A**) and (ii) donor response (**Figure 4B**) for each peptide, similar to the original RATE program. By default, the web implementation of COMBO-RATE automatically pairs the alpha and beta chains of the HLA-DP and -DQ alleles contained in the donor file, and uses a cutoff of 1 for the response. Optionally, COMBO-RATE allows filtering HLA–peptide associations based on the results of a binding prediction method found in the IEDB website. The percentile-rank is a custom metric added by the IEDB implementation of the MHC-binding prediction methods that measures the relative position of the HLA–peptide prediction within a reference set of 10,000 random sequences derived from Swiss-Prot. By default, COMBO-RATE selects the top 25%-ile predicted outcome. For each response threshold cutoff, COMBO-RATE outputs a series of files containing the RATE restrictions for each peptide separately, along with a file that compiles the results for all peptides (**Figure 4C**). Additionally, the file summary_restrictions.csv summarizes all HLA restrictions identified in the COMBO-RATE run (**Figure 4D**).

**Figure 4:**
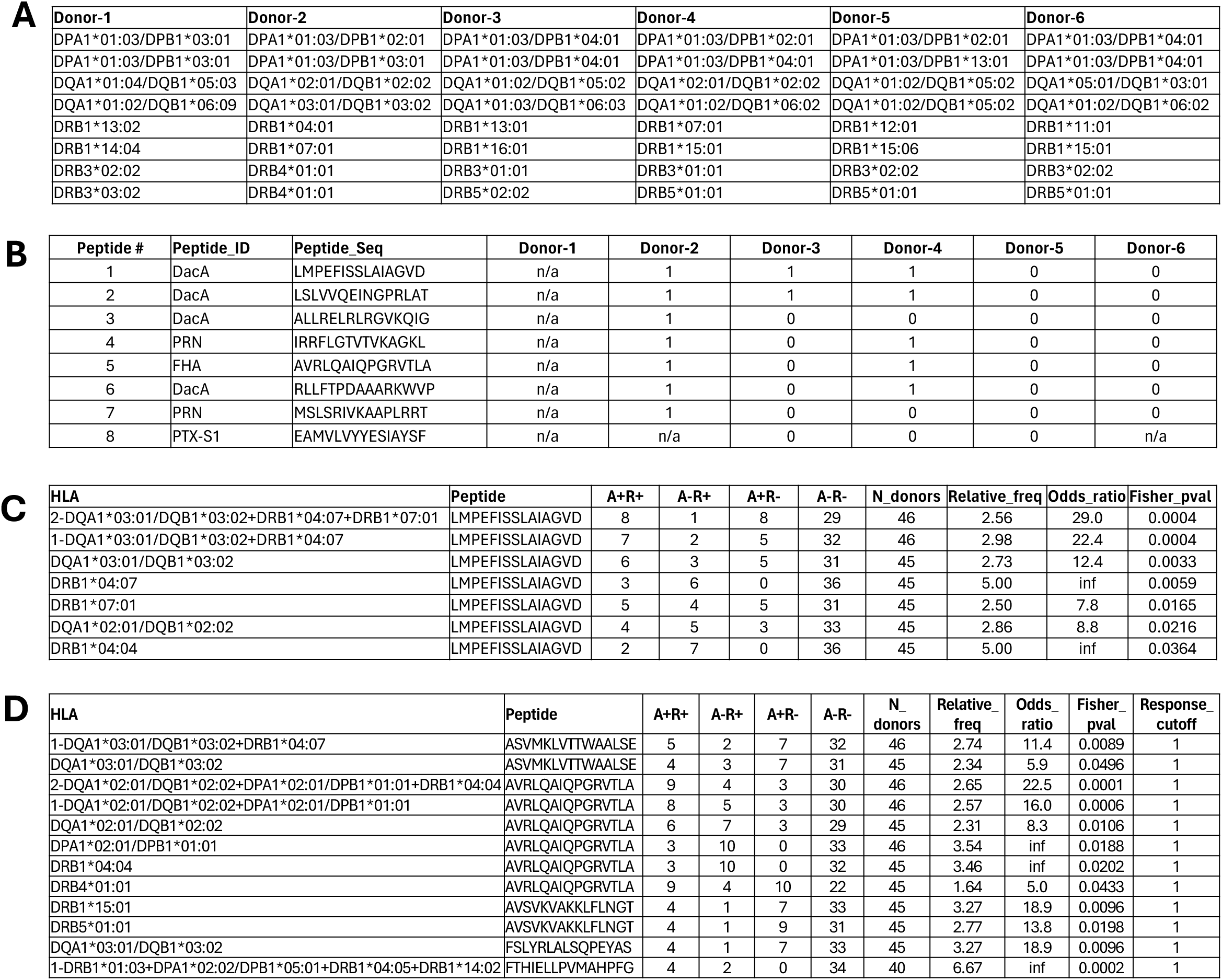
A web-based interface for community access to COMBO-RATE. **A)** Input format for the donor file in COMBO-RATE. Each column contains the allele list for a different donor identified by the column header. **B)** Input format for the response file in COMBO-RATE. Each peptide sequence is identified by a number and an ID. From the 4th column on, the response of each donor to the peptide is recorded. **C)** COMBO- RATE output file for peptide 1 (LMPEFISSLAIAGVD). Each HLA-peptide association is shown along with its respective A+/-R+/- data, number of donors, relative frequency, odds ratio, and p-value of the Fisher test. COMBO-RATE restrictions are denoted with the -1 and -2 prefix. **D)** Summary output file. For each peptide, the HLA restrictions found by COMBO-RATE are shown.

## DISCUSSION

Understanding the HLA restriction of T cell epitopes is important to characterizing immune responses in the context of infectious diseases, autoimmunity, and vaccine development. While single HLA restrictions are often assumed or reported in the literature^13,20-24^, promiscuous HLA restriction, wherein a single peptide epitope is presented by multiple distinct HLA molecules, has been recognized and studied for over three decades^11,25-29^. What remains poorly understood, however, is how often promiscuous restriction occurs at the population level, and whether recognition across multiple HLA class II molecules plays a significant role in driving epitope immunodominance. Furthermore, existing tools for inferring HLA restriction from population response data, including the RATE algorithm, were not designed to detect or report multi-allele restrictions and can be statistically confounded by them. The present study addresses both of these gaps.

Our analysis of three independent datasets spanning distinct antigen targets (*Bordetella pertussis*, Measles virus, and C9orf72), reveals that promiscuous restriction is not only common but near-universal among immunodominant CD4+ T cell epitopes. Across the 43 dominant BP epitopes tested, only 3 epitopes (7%) could be attributed to a single HLA allele present in all responders; in all remaining cases, no single HLA molecule was shared across the full responder group. Notably, this pattern was not a consequence of the computational epitope prediction strategy used to pre-select the library of BP epitopes screened^30,31^. An identical pattern of multi-allele restriction was observed in datasets derived from screening entirely unbiased overlapping peptide libraries spanning the Measles proteome and C9orf72^19,32^. The inverse correlation between the frequency of the most common HLA among responders and the total number of responders further underlines this point. Immunodominance, defined by recognition frequency of an antigen or specific epitope regions across a diverse population, is likely itself a consequence of broad HLA compatibility. While the existence of promiscuous epitopes has long been appreciated, the near-complete absence of single-allele restrictions among dominant epitopes, demonstrated here systematically across independent antigens and selection strategies, is a quantitative finding that invites reconsideration of how HLA restriction is typically characterized. It also suggests that the breadth of HLA restriction, rather than single-allele binding, may be a more informative metric when prioritizing dominant epitopes.

The RATE algorithm, available through the IEDB^15,16^, provides a statistical framework for inferring HLA restrictions from epitope response patterns in HLA-typed cohorts. By identifying alleles overrepresented among responders relative to non-responders, RATE captures restriction signals in many cases. However, when multiple HLA alleles independently restrict the same epitope, as our data suggest is the norm, single-allele association testing is systematically underpowered. This dilutes the apparent association of each individual allele and identifies the most statistically prominent restriction in many cases, but misses potential additional restrictions. To address this limitation, we developed COMBO-RATE, an iterative extension of RATE that evaluates combinations of different HLA alleles. As expected, COMBO-RATE applied to the BP dataset substantially outperformed conventional RATE. The fraction of restriction-matched donor-peptide response instances increased from 35% to 55%, or 67% when restricted to epitopes for which at least one restriction could be inferred. Restrictions were identified for 35 of 43 epitopes compared to 24 with RATE alone, and COMBO-RATE identified 64 new allele-level restrictions including 29 unique alleles undetected by single-allele analysis. Importantly, COMBO-RATE recovered restrictions for 11 epitopes where RATE returned no significant association at all, and yielded markedly improved p-values across the epitopes where both methods identified restrictions.

A key strength of the present study is the experimental validation of COMBO- RATE-inferred restrictions, which has never been comprehensively performed in prior studies attempting to infer multiple HLA class II allele restrictions^13,16,33^. Using a broadly recognized epitope across multiple donors, we confirmed that both DRB1*15:01 and DRB1*01:01 independently present the same peptide to peptide-specific CD4+ T cells, thereby validating the ability of COMBO-RATE to identify genuine biological restrictions. Importantly, this approach also identified DRB1*01:01 as an additional restricting allele, one that failed to reach statistical significance in single-allele analysis and would otherwise have been overlooked. More broadly, by prioritizing the most likely restricting alleles, COMBO-RATE can guide experimental validation efforts in future studies by narrowing down which alleles to test experimentally, reducing the number of restriction assays and HLA-transfected cell lines needed while maximizing the likelihood of identifying true restrictions.

The near-universality of promiscuous HLA restriction among dominant epitopes has direct implications for vaccine antigen selection. Because promiscuous epitopes frequently map to conserved antigen regions^25,34,35^, they offer broad HLA coverage and reduced susceptibility to mutational escape simultaneously. A small set of such epitopes can be a substitute for larger multi-epitope constructs, and when coupled to B cell targets, can ensure broad T cell activation across genetically diverse populations, addressing a key limitation of earlier single-allele-dependent vaccine strategies^36-38^. Our study could have important implications particularly for subunit vaccine designs targeting pathogens with highly diverse global populations, where broad HLA coverage across DR, DP, and DQ loci is essential. COMBO-RATE provides a means to derive these more complete restriction profiles from existing population response data, without requiring additional experimental work beyond what is already collected in standard epitope characterization studies.

To make COMBO-RATE broadly accessible, we implemented the algorithm as both a standalone Python script (https://github.com/raphaeltrevizani/comborate_shared) and a user-friendly web application (https://comborate.onrender.com). The web interface handles several practical complexities that commonly introduce errors in manual analyses, including the automated pairing of HLA-DP and HLA-DQ alpha and beta chains. Users can optionally filter HLA-peptide pairs using IEDB binding predictions, retaining only alleles with a percentile rank of 25% or better relative to a reference set of random peptides, a step that focuses the analysis on alleles with genuine peptide-binding capacity and reduces the search space for combinatorial testing. Output files summarize both per-peptide restriction assignments and a compiled cross-epitope restriction table, facilitating downstream analysis and reporting. The COMBO-RATE web tool thereby extends the reach of the RATE framework to the broader immunological community, with no programming expertise required. Although the present study focuses on HLA class II and CD4⁺ T cell responses given the available BP datasets, COMBO-RATE also supports HLA class I restriction analysis, applying binding predictions independently for each class and generating separate outputs from a single input dataset.

Several limitations merit acknowledgment. First, the statistical power of COMBO- RATE, like that of RATE, scales with cohort size and HLA diversity. For rare alleles or epitopes with few responders, combinatorial testing may not achieve significance even when a biological restriction exists, and results should be interpreted with appropriate caution. Second, COMBO-RATE also cannot infer restrictions for alleles absent from the study population, which may lead to underrepresentation of restrictions relevant in populations not captured by the tested cohort. Third, while we provide experimental validation for one epitope, the remaining COMBO-RATE-inferred restrictions are computational predictions.

In summary, while promiscuous HLA restriction has long been recognized as a feature of CD4+ T cell epitopes, we demonstrate here that it is in fact the near-universal mode of immunodominance across independent antigens and epitope discovery strategies. Importantly, COMBO-RATE directly addresses the failure of existing restriction-inference tools to reliably detect promiscuous HLA restrictions. By providing COMBO-RATE as a freely accessible tool, we further aim to support more accurate and complete HLA restriction characterization, with downstream benefits for vaccine design, population coverage, and mechanistic studies of antigen presentation.

## METHODS

### Human subjects

Blood samples were collected from healthy adult donors through the Clinical Core Facility at La Jolla Institute (LJI) in San Diego, California. The study cohort was approved by the LJI Institutional Review Board protocol # VD-214, and all participants provided informed consent. Blood was collected via venipuncture into heparin-coated collection bags. Peripheral blood mononuclear cells (PBMCs) were isolated by density-gradient centrifugation using Ficoll-Paque Plus (Lymphoprep), as preformed in previous studies^4,39,40^. Isolated PBMCs were cryopreserved in a solution containing 90% heat-inactivated fetal bovine serum (FBS; Hyclone laboratories, Logan, UT) and 10% dimethyl sulfoxide (DMSO; Gibco) and stored in liquid nitrogen until use.

### HLA Typing

Genomic DNA from PBMCs was typed at the American Society for Histocompatibility and Immunogenetics (ASHI), an accredited Institute for Immunology and Infectious Diseases (IIID), Murdoc University (Western Australia). Locus-specific, sample-indexed (multiplex identifier, MID) primers targeting polymorphic exons were used for Class II HLA typing, including HLA-DQA1/-DQB1 (exons 2-3), HLA-DRB1/3/4/5 (exon 2), and HLA-DPB1 (exon 2). Since sequencing was limited to these regions, variation outside the targeted exons cannot be excluded, and some alleles are reported as G-group assignments where phase could not be resolved. Amplicons (up to 96 MIDs per run) were quantified, pooled equimolarly, and prepared using NEBNext Ultra II library preparation kit. Library concentration was measured using a Quantus Fluorometer (Promega), and fragment size distribution was assessed with High Sensitivity D1000 ScreenTape on an Agilent 2200 TapeStation. Sequencing was performed on an illumina MiSeq (v3, 600-cycle, 2x300 bp). Reads were demultiplexed (by MID barcodes), trimmed, quality-filtered, and merged prior to allele assignment using the IIID HLA Analysis Suite with the IMGT/HLA database as a reference. Quality control metrics, including read depth, allele balance, and contamination checks, were monitored using the IIID Laboratory Information Management System (LIMS). This workflow does not type HLA-DPA1. Therefore, DP restrictions shown as DPA1/DPB1 pairs, are derived from inferred pairing rather than direct genotyping.

### Validation of inferred restrictions using HLA II-transfected cell lines

Donors responding to P23 (**Table S3**) were selected based on high resolution HLA II genotyping to express either HLA-DRB1*15:01 or HLA-DRB1*01:01, but not both. Donors expressing DRB1*08:02 or DRB1*14:02 were excluded to minimize ambiguity in restriction assignment. Candidate peptide P23 (1mg) was synthesized at >99% purity (TC Peptide) and reconstituted to 20mg/mL in DMSO. PBMCs were thawed and cultured in HR5 (RPMI + 5% human serum + 1% Pen-Strep + 1% Glutamax) medium. Cells were plated at 12x10^6^ cells/well in 6-well plates and stimulated with 1ug/mL of peptide. Cultures were maintained at 37C, 5% CO_2_ for 14-days, with recombinant IL-2 (10 U/mL) added every 3-4 days. To validate HLA II restricted antigen presentation, DAP.3 fibroblast expressing either DRB1*15:01 (L466.1) or DRB1*01:01 (L554.3) were used as APCs. The parental DAP.3 cell line lacking HLA-DR expressing was utilized as a negative control (**Sup. Figure 1**). Cell lines were maintained in R10 fibroblast media (RPMI + 1% non-essential amino acids + 1% Sodium Pyruvate + 10% FBS + 1% Glutamax +1% Pen-Strep) supplemented with G418 (Geneticin). HLA II expression was induced by treatment with sodium butyrate (0.1mg/mL) for 24 hours prior to use. HLA-DR expression was validated by flow cytometry. Briefly, cells were detached using 0.05% trypsin-EDTA (1x) solution (Gibco) washed twice with 1x PBS and incubated with anti-HLA-DR (PE) antibody (clone L243; BioLegend) and fixable viability dye eFlour506 (eBioscience) for 40 minutes at 4C. Stained APC samples were acquired on a ZE5 cell analyzer (Bio-Rad laboratories, Hercules, CA) and analyzed with FlowJo software (Tree Star, Ashland, OR) (**Sup. Figure 1**). FluroSpot plates (Mabtech) were coated overnight at 4°C with a mouse anti-human IFN-γ (clone 1-D1K, Mabtech) antibody. APC cell lines were detached using 0.05% trypsin-EDTA (1x) solution (Gibco), washed and pulsed with 1ug/mL of peptide and incubated for 1 hour at 37C, 5% CO_2_. Cells were washed five times with HR5 media to remove excess peptide. Unpulsed cells were processed in parallel and used as negative controls.

The IFN-γ FluoroSpot assay was used to quantify antigen-specific T cell responses. PBMCs (50,000 cell/well) from each donor were co-cultured with P23-pulsed or unpulsed APC (12,500 cell/well) as follows: 1) PBMCs + L466.1 (DRB1*15:01), 2) PBMCs + L554.1 (DRB1*01:01), 3) PBMCs + Parental on coated FluoroSpot plates. Additionally, PBMCs were stimulated with peptide only (10ug/mL) and PHA (10ug/mL) as positive controls and DMSO as a negative control. All conditions were tested in triplicate. Plates were incubated for 20-24 hours at 37C, 5% CO_2_. After incubation, cells were removed, and membranes were washed. An IFN-γ (7-B6-1-FS-BAM) prepared in PBS with 0.1% bovine serum albumin (BSA) was added and incubated for 2 hrs at room temperature. Membranes were then washed, and plates were incubated with an anti-BAM-490 (Mabtech) secondary antibody for 1 hr at room temperature. Plates were subsequently washed, incubated with fluorescence enhancer (Mabtech) for 15 minutes, and air-dried befor reading. Spots were read and counted using the Mabtech IRIS system Criteria for positivity were as follows: 1) A minimum net response of >20 SFC per 10^6^ PBMCs after subtracting DMSO background control. 2) The T cell reactivity values measured in triplicate in response to background need to be significantly higher (p<0.005), as assessed by Welch’s t-test and/or Poisson test. 3) T-cell reactivity observed needed to reach a stimulation index (SI) >2 (e.g., have a magnitude of at least 2-fold higher than background).

### RATE analysis

Briefly, RATE^15,16^ takes two files as input. The first details the HLA alleles for each donor (**Figure 4A**) and the second contains the measures of the responses for each peptide and donor combination (**Figure 4B**), both sharing the same output as COMBO-RATE (see below). RATE counts the occurrences of allele positive donors (A+), responders (R+), allele negative donors (A-) and non-responders (R-) and, based on these values, estimates the odds ratio (OR), the relative frequency (RF), and runs a Fisher exact test. Briefly, in OR are odds where the numerator represents concordant pairs (both positive or both negative) and the denominator represents discordant pairs (one positive, one negative) according to:

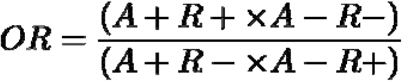

RF is estimated as the ratio of two proportions: the proportion of responders among allele-positive individuals and the overall proportion of responders in the full cohort. Specifically, the first term is the fraction of allele-positive individuals who are responders (A+R+/A+), and the second term is the fraction of all individuals who are responders (R+/total donors). The RF is then defined as the ratio of these two quantities, according to:

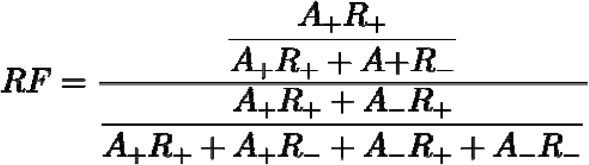

Finally, the A+R+, A+R-, A-R+, A-R- are used to make a contingency table and analyzed by a Fisher’s exact test.

### COMBO-RATE

COMBO-RATE iteratively applies RATE while progressively grouping HLA alleles. The first iteration is an ordinary RATE run. In the second iteration, it selects the two HLAs with the lowest Fisher’s exact test p-values with RF>1, and defines a set S = {p_1_, …, p_x_} where p_1_ is the HLA with the lowest p-value and p_x_ is the HLA with the second lowest p-value. All HLAs in S are treated as a single combined variable (denoted S_HLA_) across donors, and a new Fisher’s test p-value is computed. If the p-value for S_HLA_ is lower that p_1_, the set S is expanded to include the HLAs associated with the next lowest p-value (S = {p_1_, …, p_x_, p_y_, …, p_z_}, where p_y_ … p_z_ are the HLA alleles with the third lowest p-value. The algorithm continues as the p-value for S lowers and stops otherwise.

If desired, the web implementation of COMBO-RATE can also pair the alpha and beta chains of the HLA-DP and -DQ alleles contained in the donor file. All four potential combination per HLA-DP and -DQ alleles will be created. In addition, a file containing the predicted binding affinity between all HLAs expressed by tested donors and all tested peptides, determined by NetMHCIIpan 4.1 EL prediction method, can be included in the COMBO-RATE analysis to ensure that only HLAs likely to bind to a peptide are included in the analysis. HLAs without predicted binding capacity will not be included in the analysis.

## Acknowledgments

We wish to acknowledge all the subjects for their participation and for donating their blood and time for this study. This study was funded by the National Institutes of Allergy and Infectious Diseases (NIAID) Human Immunology Project Consortium (HIPC) Grant # U19 AI118626, the Cooperative Centers for Human Immunology (CCHI) Grant # U19 AI142742, the National Institutes of Health (NIH) Epitope Discovery Program contract number 75N93024C00056, and the National Institutes of Health (NIH) Ruth L. Kirschstein National Research Service Award (NRSA) T32AI125179.

## Author Contributions

J.N.M., R.d.S.A., A.S. and R.T. designed the study. R.T. developed the COMBO-RATE pipeline, J.N.M., A.A., and A.S. and performed experiments and data analysis. A.G. and E.J. provided training datasets that contributed to the study design. J.N.M., R.d.S.A., A.S. and R.T. wrote the manuscript. All authors read and approved the manuscript.

## Supplementary Figure Legends

**Supplementary Figure 1.**
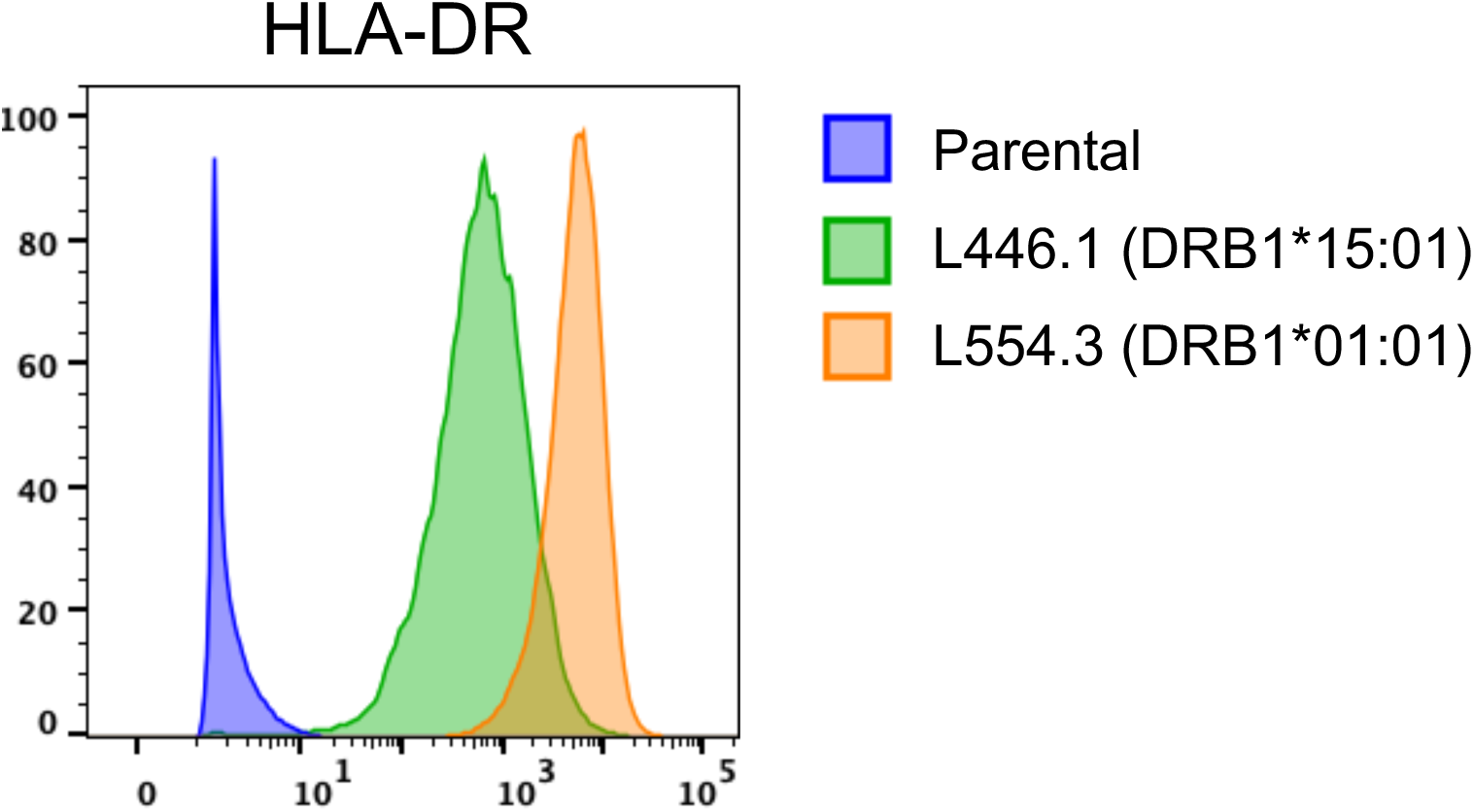
HLA-DR expression levels of single HLA II transfected cell lines. HLA-DR expression of DAP.3 fibroblast L446.1 (DRB1*15:01) and L554.3 (DRB1*01:01) cell lines compared to parental cell line expressing no human HLA II molecules.

**Supplementary Figure 2.**
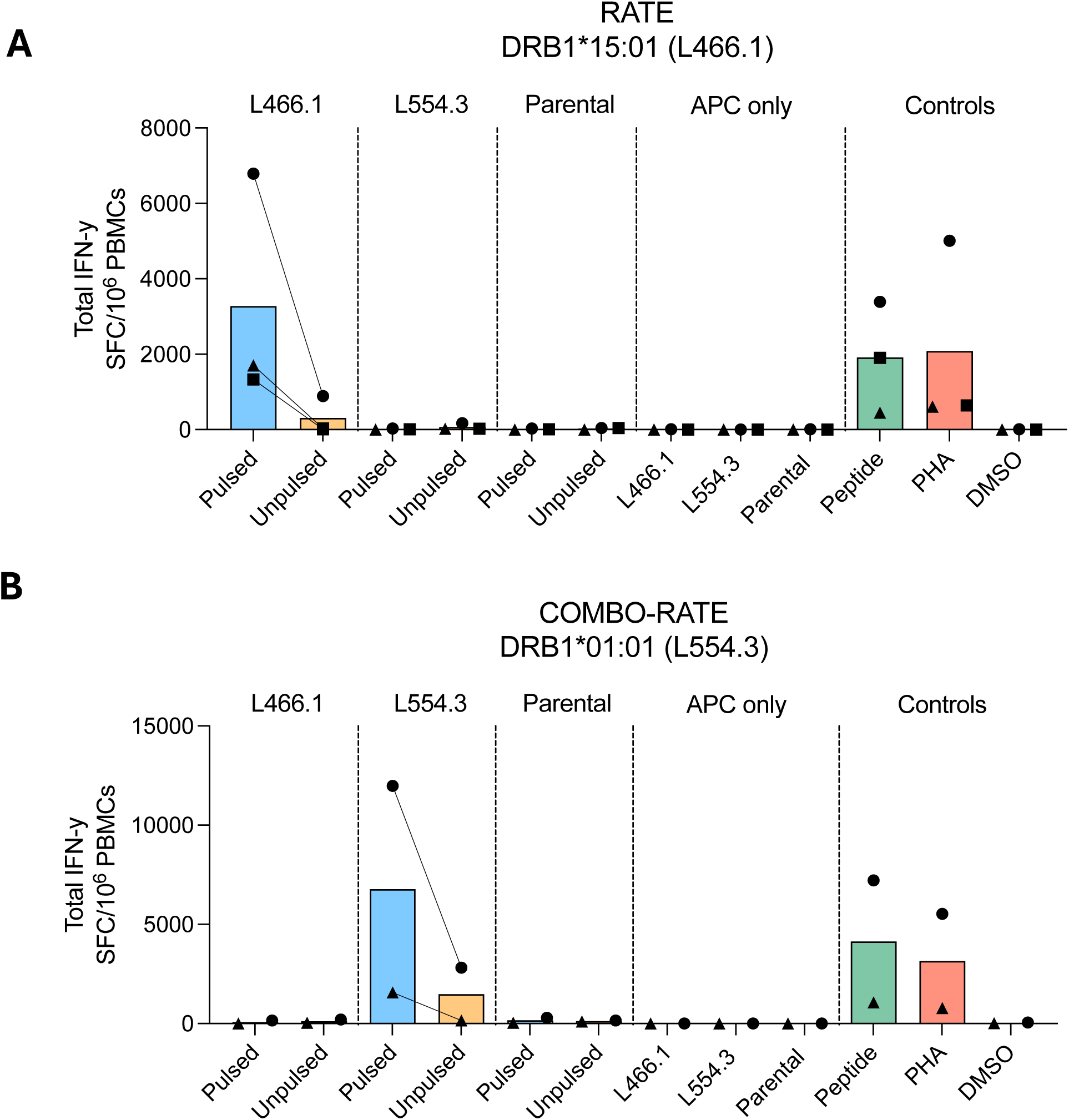
Raw data of experimental validation of RATE and ComboRATE restrictions. **A)** Total IFN-y levels (spot forming cell (SFC)/10^6^ PBMCs) of RATE-associated DRB1*15:01 allele donors (n=3) when co-cultured with P23-pulsed and unpulsed antigen presenting cells (APCs): L466.1 (DRB1*15:01), L554.3 (DRB1*01:01), and parental APC lines. **B)** Total IFN-y levels (SFC/10^6^ PBMCs) of COMBO-RATE-associated DRB1*01:01 allele donors (n=2) when co-cultured with P23- pulsed and unpulsed APCs: L466.1 (DRB1*15:01), L554.3 (DRB1*01:01), and parental APC lines. For both assays, unpulsed APC lines only were used as negative controls. PBMCs stimulated with peptide (10ug/mL) and PHA (10ug/mL) were used as positive controls and DMSO as a negative control. Each tested donor is represented by a unique shape.

**Table S3:**
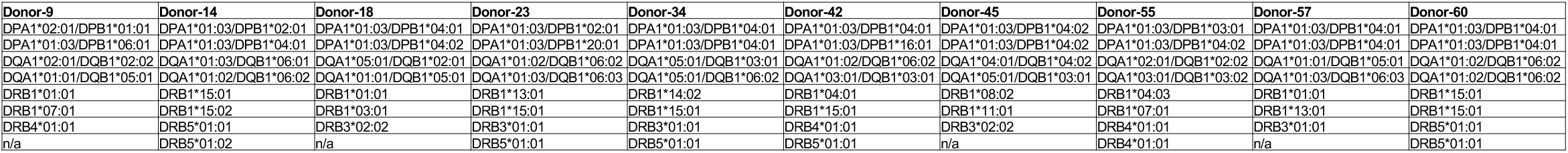
HLA type of 10 individuals responding to epitope P23.

